# ChemOrigins: A community curated database for origins of life chemistry

**DOI:** 10.1101/2025.06.10.658922

**Authors:** Bruno Cuevas-Zuviria, Zachary Adam, Daniel Cove, Betül Kaçar

## Abstract

The origin of life is one of the most compelling questions in science. While experimental prebiotic chemistry has produced a wide range of reactions and plausible pathways, the resulting data remain fragmented across numerous publications and disciplinary journals. Here, we introduce *ChemOrigins*, an open-access, community-curated knowledge graph that organizes experimentally supported prebiotic reactions. By representing molecules, reactions, conditions, and literature sources as interconnected nodes, ChemOrigins enables modular grouping of reactions and supports complex, query-driven exploration via a graph database architecture. We demonstrate the utility of this framework through text-based searches, reaction network expansions, and the interactive visualization of user-annotated chemical modules. Unlike generative models, ChemOrigins prioritizes curated, evidence-based content and fosters community contributions through expert annotations and a user-friendly interface. As a structured resource, ChemOrigins is designed to complement existing chemical databases and serve as a foundation for computational, educational, and theoretical research in the origins-of-life field.

## Introduction

Understanding how life emerged remains one of science’s most profound challenges. Researchers aiming to bridge the synthesis of biomolecules with the emergence of complex systems must navigate a vast and rapidly growing body of prebiotic chemistry literature [1,2]. Yet, navigating this landscape is inherently difficult: proposed pathways are dispersed across numerous specialized and generalist journals, and interpreting them often requires expertise in organic chemistry, geochemistry, and the merits and weaknesses of diverse origins-of-life conceptual models.

By contrast, neighboring disciplines such as gene regulation, mineral formation, and atmospheric chemistry have increasingly adopted systematic approaches to organize and interconnect the processes they study. These efforts have not only improved data accessibility across disciplines but have also enabled advanced applications such as database integration, cross-referencing and machine learning driven analyses [3–5]. Prebiotic chemistry could similarly benefit from a structured, comprehensive resource: one that systematically compiles key theories, reactions, and experiments in a format that is both machine-readible and user-friendly. Such a platform could reduce entry barriers, bridge disparate theoretical models, and open new avenues for research.

To address this need, here we introduce an open-source, community-driven database specifically designed to catalog findings in prebiotic chemistry using an interoperable format. As a foundation, we manually curated and annotated over 300 experimentally supported prebiotic reactions to seed the database. This platform is openly accessible and designed to support both human exploration and machine driven programmatic access, available at http://chemorigins.bact.wisc.edu.

## Results

Developing ChemOrigins required designing and implementing protocols that span the entire data life cycle, from initial data curation to end-user website interface. **Figure 1** illustrates the data life cycle: curators generate annotations based on either their own experimental work or from the relevant literature. These annotations are then converted and uploaded to a graph database, which enables complex queries. Finally, a web interface allows access to this database through multiple search tools.

**Figure 1.**
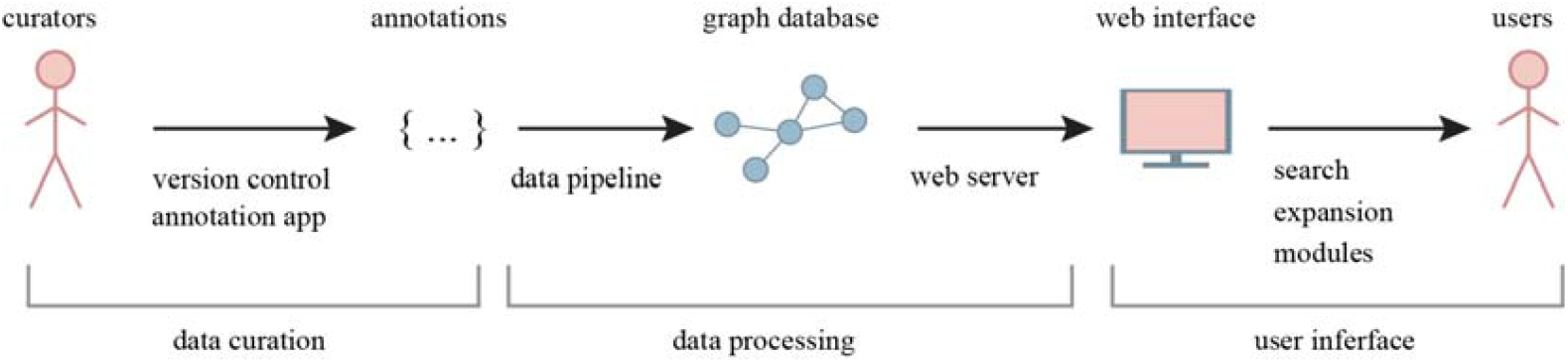
ChemOrigins data life cycle: from curators, who create the original annotations, to end users through the graph database and the web interface. In gray, we included some of the components of ChemOrigins, such as the data processing pipelines or the version control system that enables collaborative curation of prebiotic chemistry.

### Data: Annotations and data curation

Systematizing prebiotic chemistry is a long-term endeavor that depends on continuous community contributions. However, the high cost of employing dedicated scientific curators presents a challenge. To address this, we adopt a community-based curation model inspired by successful precedents, such as the Open Reaction Database [6], but specifically tailored to the unique demands of prebiotic chemistry.

The fundamental unit of ChemOrigins is the annotation (**Figure 2*a***), a document generated by curators that describes a specific piece of information from the literature. Ideally, each annotation is a computer-readable declaration describing a reaction, molecule, agent, or related entity. In this initial version, we focus on chemical reaction annotations, which describe the conversion from reactants into products as experimentally reported in scientific publications. To ensure both traceability and machine readability, we require users to provide two indispensable attributes: the DOI of the original source and the reaction in SMILES format. Additional optional fields include reaction conditions, agents, comments, and more.

**Figure 2.**
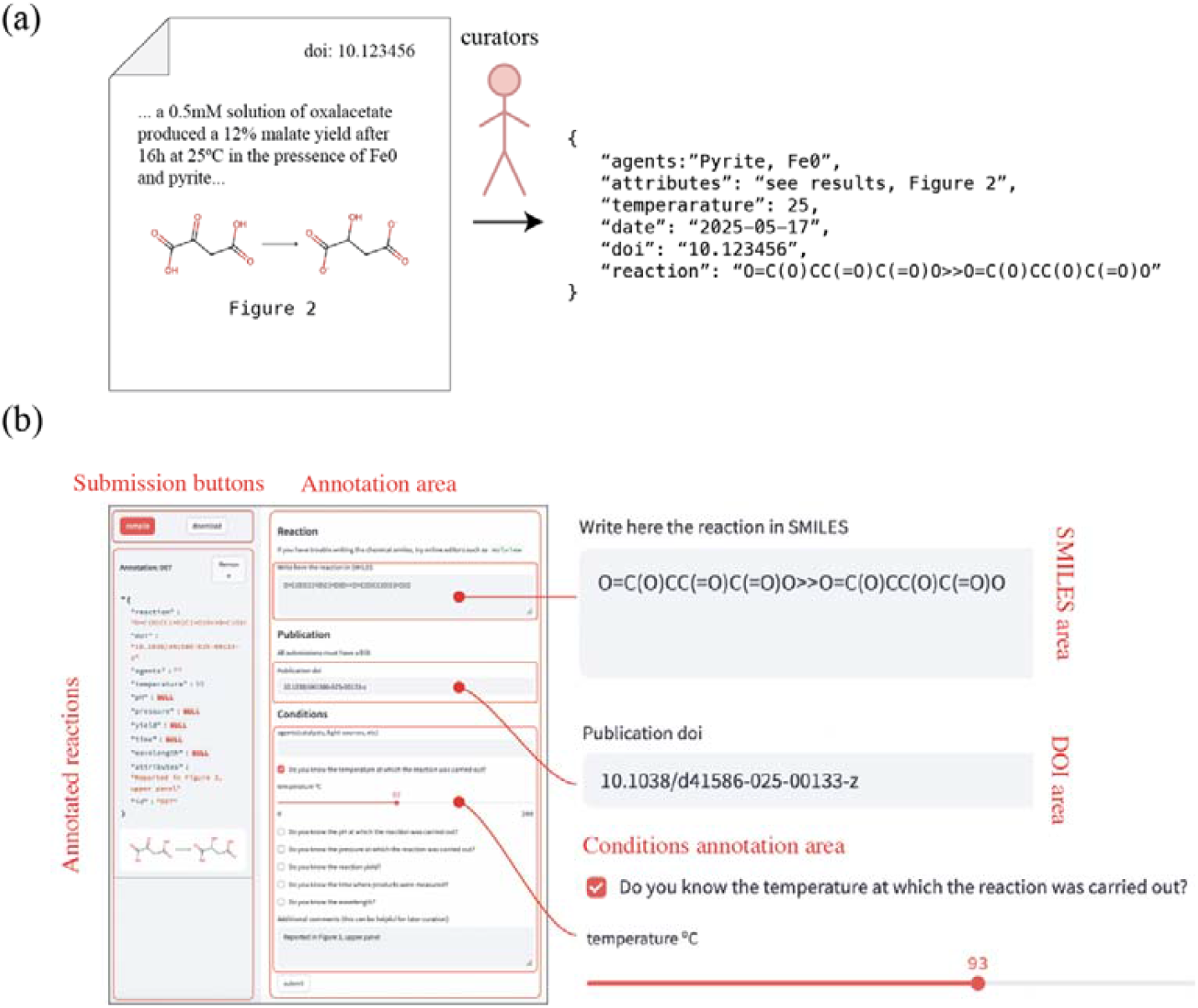
Explanation of the curation process. **(*a***) Users find an interesting piece of literature that describes a given set of conditions. Based on that, they can upload an annotation, which must follow specific rules to be processed by the data processing pipeline. **(*b*)** An annotation app allows users to create their own entries and upload them without requiring them to familiarize themselves with the complexities of version control systems.

Annotations are collected and maintained in an online repository using a version control system (git). This approach offers three main advantages. First, it preserves a complete history of edits, allowing any previous version to be restored if needed. Second, it supports collaborations: users can create branches of the main repository to submit new data, which are then reviewed and merged by database administrators. Third, it ensures the open access, allowing others to build upon this dataset for future research and applications. The repository is publicly hosted at https://github.com/brunocuevas/chemorigins-data.

Although the usage of version control systems such as git is becoming more widespread, we recognize that it may impose a technical barrier for many intended users. To mitigate this, we developed an annotation app that enables users to create and submit annotations directly (**Figure 2*b***), through a guided interface. The app presents a user-friendly form where each field relevant to the reaction is filled in directly. The app takes charge of the cumbersome version control operations in the background, liberating the user from these tasks and making participation more accessible.

### Backend: Prebiotic knowledge graph

At the core of the ChemOrigins database lies a prebiotic knowledge graph, which underpins complex queries and enables advanced analytical capabilities. In our implementation, the graph structure (**Figure 3*a***) consists of nodes and edges. Nodes represent distinct entities, such as molecules, chemical reactions, experimental conditions, and literature sources. Edges encode the relationships between these entities. This design offers flexible and extensible data integration, reflecting the inherently interdisciplinary nature of prebiotic chemistry.

**Figure 3.**
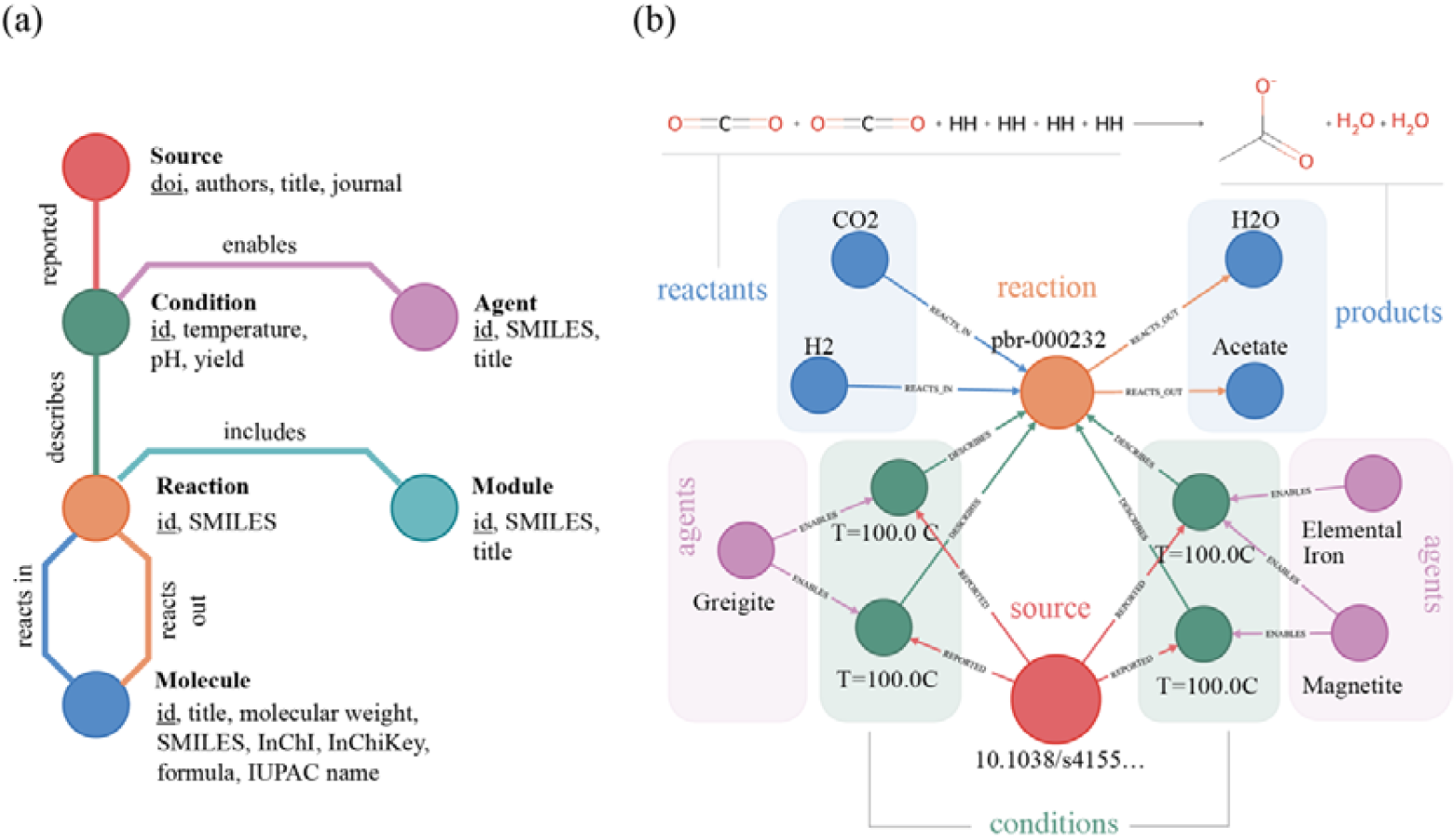
Representation of the schema employed in ChemOrigins. **(*a*)** Graph highlighting the different kinds of entities that we might find represented in ChemOrigins, together with some of their attributes (e.g., temperature, SMILES, etc) and the relationships between them. **(*b*)** Graph representation of the data associated with a given reaction (shown at the top of the figure), including substrate and product molecules (blue), the conditions at which those reactions took place (in this case, we highlight the temperature), the agents that allowed that reaction (in this case, different iron associated minerals) and finally, the source of the data, which is identified by its DOI.

To illustrate the capabilities of this structure, **Figure 3*b*** shows a representative reaction connecting specific reactants to products, all reported under multiple experimental conditions within a single publication. In this example, we linked an agent (neutral iron) to the reaction via a condition node, reflecting its role as a catalyst. Importantly, the graph architecture accommodates such context-dependent roles: the same reaction may be associated with different catalysts under varying conditions, each captured as a distinct connection. The resulting network can be further organized into higher-level groupings when subsets of reactions collectively describe, for example, a plausible prebiotic synthesis route to a compound of interest. To capture this, we introduce “module” nodes that connect and label related reactions, enabling users to identify and analyze relevant sub-networks within the larger reaction space (see next sections, *Frontend: powering prebiotic chemistry search*, and *New insights from systematized chemistry*).

One key advantage of using a graph-based representation is the conceptual alignment with the nature of the chemical process. In chemical systems, species participate in reactions that lead to new compounds, forming intricate networks, some of which have been hypothesized to play critical roles in the emergence of life [7–18]. A graph-based model captures these relationships in a way that mirrors their real-world behavior relatively better, offering a more intuitive and flexible alternative to traditional tabular representations. This approach also simplifies the formulation of complex queries that involve traversing reaction networks. Tasks that would be cumbersome or inefficient in conventional database formats become straightforward within a graph framework.

For example, Arya et al. [11] employed a similar representation and database engine to identify autocatalytic motifs in rule-generated chemical reaction networks, demonstrating the analytical power of this method. In **Figure S1**, we present examples of Cypher queries that can be performed using this representation. **Figure S1a**, shows how to find all molecules involved in reactions described in a specific paper [19]; **Figure S1*b*** identifies all reactions catalyzed by a particular agent, and **Figure S1*c*** expands all reactions involving both formaldehyde and cyanide simultaneously, along with their associated catalysts. These examples underscore the potential of the knowledge graph to support detailed exploration and hypothesis generation in prebiotic chemistry.

### Frontend: powering prebiotic chemistry search

The data curated by the community are available for users through either a web portal or a REST API. In the first case, users can access https://chemorigins.bact.wisc.edu with their browsers and make direct queries to the website (**Figure 4*a*)**. In the second case, applications can request data from ChemOrigins by using the REST endpoint (e.g., https://chemorigins.bact.wisc.edu/api/reactions/pbr-000232). More detailed information about the REST API can be found in the website documentation. Here, we focus on the description of the web interface.

**Figure 4.**
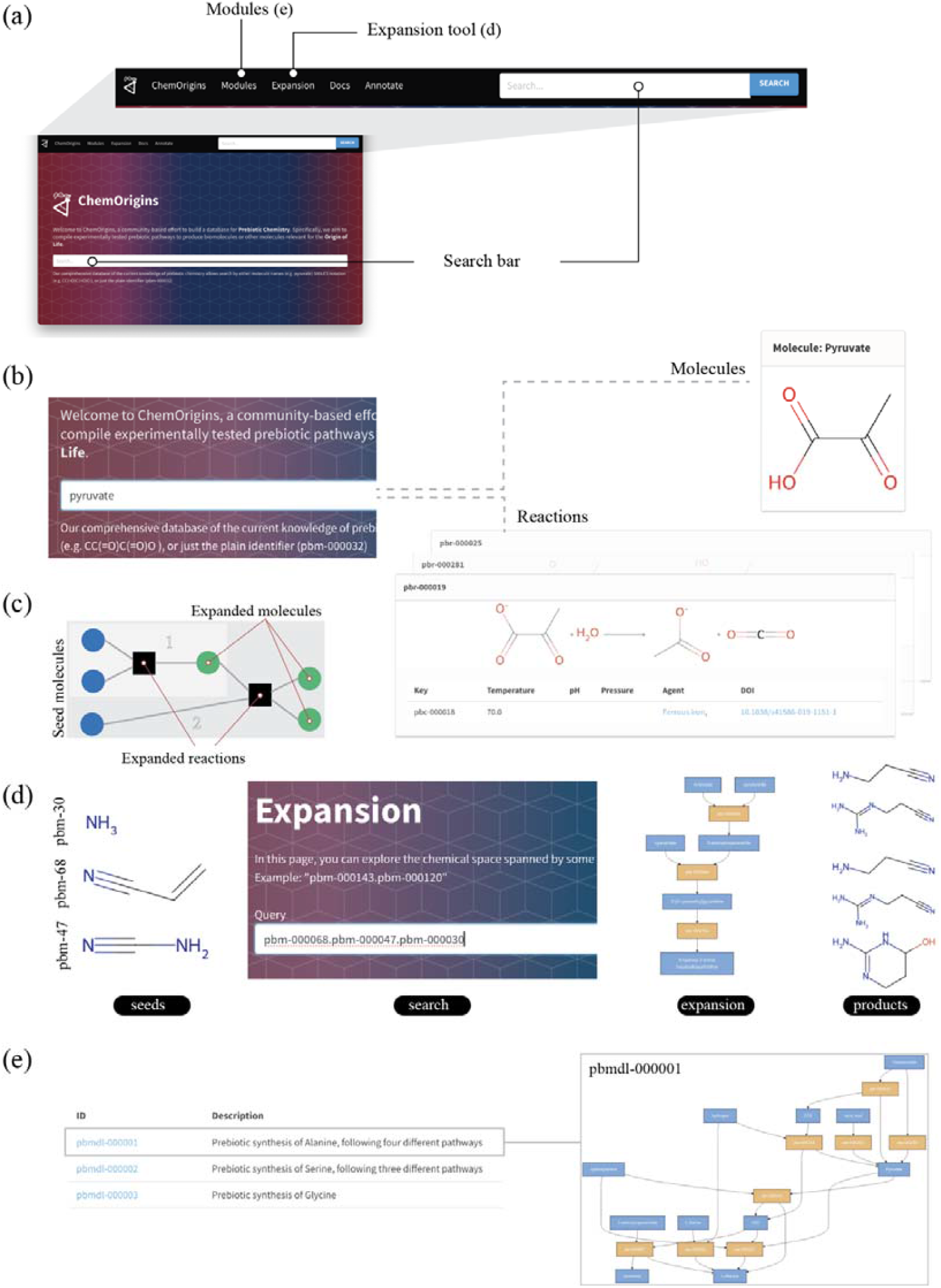
ChemOrigins web interface. (***a*)** Location in the landing page of the three main methods to interact with the database: search, expansion and modules. (**b**) Search mode: Introducing a text query, such as the molecule name, its SMILES, or its identifier, leads to the collection of molecules and reactions that match the query. (**c**) Example of network expansion: the method identifies a collection of reactions and molecules spanning from a set of seed molecules. (**d**) Example of expansion based on a set of three molecules, illustrating the result as a diagram and as a set of molecules and reactions found at the expansion. (**e**) Modules are listed with a short description of their role. Opening a module leads to a page that includes the molecules, reactions, and a diagram, as shown in the figure.

Querying the database may be the most critical aspect of the user interface; ideally, ChemOrigins should enable the retrieval of information that would otherwise require hours of reading in just a few seconds. To support this, we have implemented three modes of interaction: i) regular search, ii) expansions, and iii) modules. The first mode (**Figure 4*b***) is the most intuitive example: the user types in a search bar and retrieves a list of related items: molecules, reactions, etc. The second mode is more specific to prebiotic chemistry. Chemical reaction network expansions (**Figure 4*c***) allow us to obtain the chemical space (i.e., a set of molecules and reactions) spanned by a defined set of initial substrates. These expansions have been successfully applied in prior origins-of-life studies to investigate proto-metabolisms [12,14,15,20]. While such applications are particularly relevant to the field, network expansions also provide a powerful tool for database exploration: by supplying a set of starting molecules, users can retrieve all downstream reactions and molecules (**Figure 4*d***).

Finally, we implemented a feature inspired by other databases, such as the Kyoto Encyclopedia of Genes and Genomes (KEGG) [21]: categorization of reactions into modules. These modules are curated by users through a straightforward annotation scheme (simply, a list of reaction identifiers), and allow the visualization of reaction sets relevant to particular contexts (for example, reactions leading to the production of alanine, **Figure 4*e***). In addition to these search tools, ChemOrigins allows users to browse information on molecules, reactions, agents, and publications by navigating the hyperlinks available and downloading formatted data in JSON.

### New insights from systematized prebiotic chemistry

Systematizing prebiotic chemistry may prove valuable in two ways. First, it facilitates education by allowing prebiotic chemists to easily explore prebiotic pathways. Second, it may lead to new insights, much like analyses of existing metabolic databases have yielded novel hypotheses about the origins and early life [12,15]. In the following paragraphs, we illustrate some of these possibilities through curated modules.

The first case (**Figure 5*a***) illustrates how ChemOrigins can help users explore entire chemical pathways leading to a target compound. We manually compiled a module (pbmdl-000001), representing reactions that produce Alanine from various starting points. This module highlights three distinct pathways: synthesis from pyruvate, formation via Strecker synthesis [22], and reduction of serine in the presence of molecular hydrogen. Each pathway corresponds to different environmental scenarios: synthesis from pyruvate is relevant to hydrothermal vent-like settings [23], Strecker synthesis may dominate in surface environments [24], and the third route underscores the potential for molecular interconversion in plausible prebiotic scenarios.

**Figure 5.**
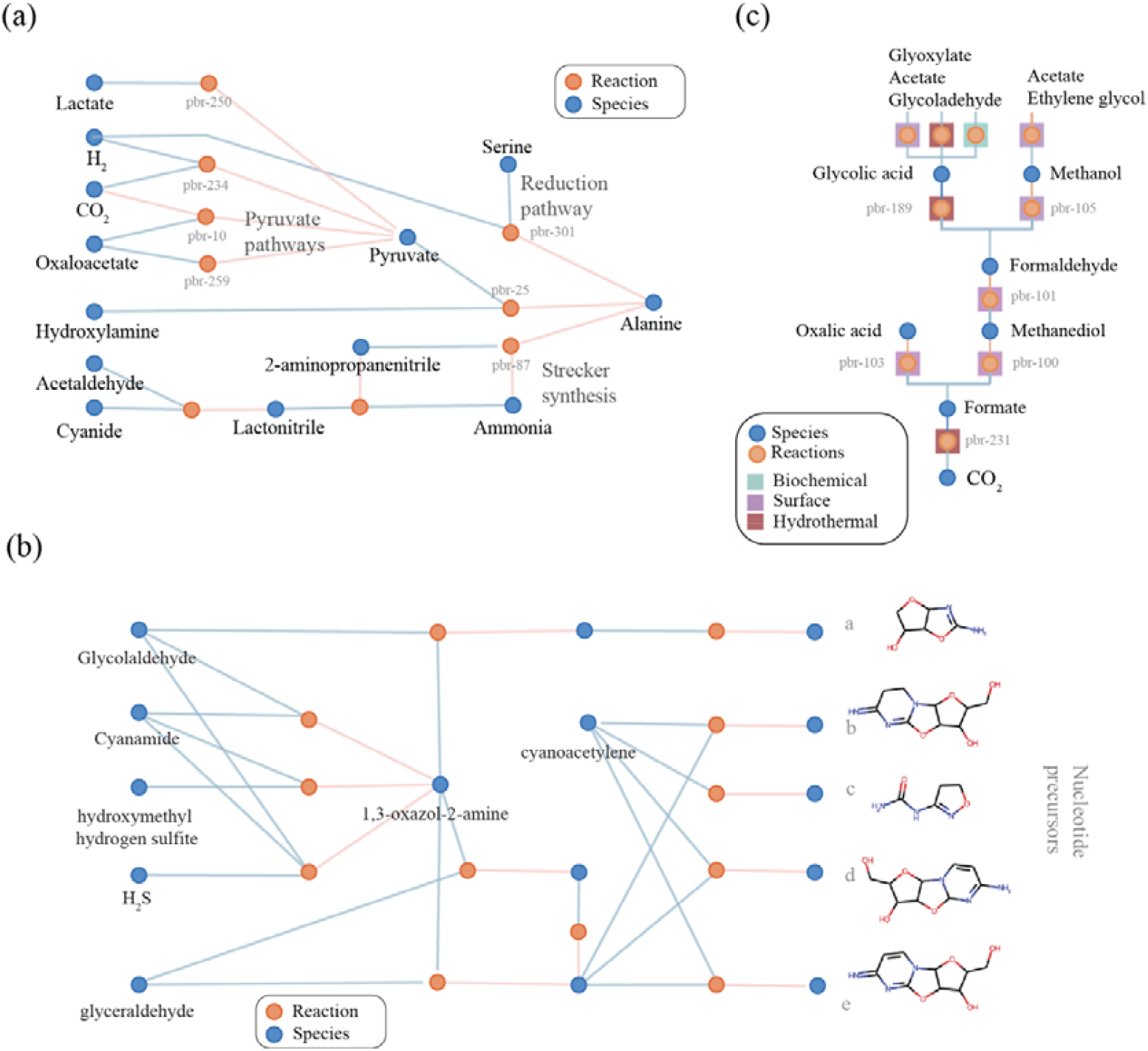
Example visualizations of ChemOrigins data. **(*a*)** Pathways leading to alanine in our database. We find three different pathways: a pyruvate-based pathway, a Strecker synthesis pathway, and a reduction pathway from serine. **(*b*)** Pathways involving 1,3-oxazol-2-amine. This compound is an essential intermediate in the synthesis of nucleotides. **(*c*)** Possible prebiotic pathway enabling carbon fixation through a combination of environments.

**Figure 5*b*** presents the reaction network centered on 1-3-oxazole-2-amine (pbmdl-000011), an essential intermediate in the Powell-Sutherland pathways for nucleotide synthesis [25]. This molecule has been synthesized under different conditions using similar precursors [7,24], and subsequent reactions enabled the formation of nucleotides. In cases (a) through (e), reactions with cyanoacetylene or glyceraldehyde generated structurally similar intermediates. When reacted in a phosphate buffer with urea, some of these intermediates lead to the formation of cyclic nucleotides [25].

The modules discussed so far are centered on specific biomolecules, but the ChemOrigins framework supports broader strategies. Modules can be defined using any grouping criteria, such as mechanistic similarity, or environmental context. **Figure 5*c*** illustrates this flexibility with a network expansion originating from carbon dioxide (pbmdl-000010). By combining reactions from multiple geochemical conditions, we were able to trace a plausible sequence leading to more complex molecules, including formate, methanediol, formaldehyde, glycolate, and methanol. Although there is no experimental confirmation that currently supports the feasibility of this transformation as a one-pot process, mechanisms that reduce spatial separation or mitigate diffusion barriers [26] could, in principle, permit such a pathway under prebiotic conditions.

## Discussion

Since the time of Carl von Linné in the eighteenth century, naturalists have sought to classify and systematize biological diversity. This effort led to a shared language that enabled communication across disciplinary and geographical boundaries. The rise of molecular biology revealed classical taxonomic definitions, such as species, can break down when applied to microorganisms. A similar challenge confronts chemistry: although biochemical systems have been codified through resources like KEGG, this reflects only a narrow slice of a vastly broader space of organic chemistry. The portion of known biochemical reactions and molecules is minuscule compared to the theoretical universe of possible organic transformations [27–29]. Estimates suggest that as many as 10 𥨐 □ distinct reactions could occur on Earth’s surface under natural conditions [30]. While systematizing all of chemistry is clearly infeasible, and arguably unnecessary, focused efforts in subfields such as prebiotic chemistry is both valuable and tractable.

Recent studies have explored large-scale, algorithmically generated prebiotic chemical networks using reaction rules [11,31]. Others have introduced novel computational frameworks, such as blockchain-based algorithms, to characterize complex reaction cycles [32]. While powerful for expanding chemical space and hypothesis generation, these methods often lack means to assess the geochemical plausibility of the included reactions. In contrast, ChemOrigins aims to curate and systematize experimentally supported hypotheses, complementing purely generative approaches. Rather than seeking to catalog all conceivable reactions, ChemOrigins focuses on mechanisms and contexts that promote meaningful chemical reactivity, contributing to the organization of origins of life knowledge. Importantly, ChemOrigins interoperate with existing databases, contributing a curated, hypothesis-driven and community-vetted dataset focused on the origins of life.

The rapid emergence of large language models (LLMs), such as GPT [33], may transform how we process scientific literature. These tools offer unprecedented capabilities to extract, summarize and structure information. Their utility has been demonstrated in domains such as metal-organic framework synthesis [34,35]. In theory, one could imagine bypassing curated databases entirely in favor of full-text AI queries. However, such an approach remains premature. First, systematization is not merely extraction; it requires scientific judgment, context and consensus. Prebiotic chemistry involves competing hypotheses and conflicting results that demand careful scholarly curation. Second, LLMs are sensitive to training data biases, and may struggle with domain-specific nuance, particularly given the relative scarcity of prebiotic literature compared to broader organic chemistry [36]. For these reasons, a curated, debated, and community-sourced dataset remains a necessary foundation, before embracing completely AI-driven discovery.

ChemOrigins provides this foundation: a trusted, evolving platform for organizing knowledge, refineming hypotheses, and uniting the community in the shared goal of understanding life’s chemical origins.

## Methods

### Community-based curation

The core of our systematization effort is building a database of the chemical reactions and their conditions. We seek a system that enables a community of researchers to self-organize and collaborate while controlling the quality and scope of the data included in the database. Open-source software provides an excellent example of this kind of effort. We divide the project participants into three groups: curators, moderators, and end-users. Curators are users who propose new data from either reviewing the literature or from their experiments. They also participate in discussions with other curators. Moderators have control of the repository discussions and validate the format and content of the proposed data. Finally, users extract information from the database.

### Annotation tool

ChemOrigins works through user-based annotations provided through version control techniques used in Open-Source software development. This means that users can download the annotations by cloning a repository, creating a new branch, committing their contributions, and pushing it to the online repository, where the moderators will merge the content with the main branch, which is periodically uploaded to the online DB service. However, requiring users to be familiar with open-source software development is an entry barrier that will prevent the community of annotators from growing. Therefore, we have built an app that allows users to produce annotations without necessarily understanding the process running behind. The app was implemented using Streamlit.

### Data processing pipeline and backend

Reaction annotation occurs by writing JSON files indicating at least two mandatory fields: reaction SMILES, publication DOI. Those annotations are uploaded into a graph-based Neo4j database through a Python-based framework, which allows storing additional reaction information as a multipartite network, enabling complex queries otherwise constrained on SQL-based databases. A parallel approach to categorizing extensive collections of synthetic prebiotic reactions has also used this kind of database [11]. The molecules of our database have been cross-referenced with PubChem and KEGG entries when possible.

We have manually reviewed the literature, generating a collection of documents that contain raw data about chemical reactivity: the reaction in SMILES format, the conditions under which each reaction was performed, and the DOI of the corresponding publication. From these annotations, we can generate a knowledge graph in which molecules are joined by reactions and where literature sources are joined with reactions through the conditions of each experiment (see **Figure 3A**).

### Frontend

We have implemented the REST API and the website front end using Flask, the schematic diagrams using Mermaid, and the molecular diagrams using rdkit Python library.

## Acknowledgments

We thank Tyma Sokolskyi and Zheng Peng for their occasional discussions that contributed to the development of this work, and Evrim Fer along with the members of the Kacar Laboratory for their valuable feedback.

## Author contributions

**Bruno Cuevas-Zuviría**: conceptualization, Writing-Original Draft, Data Curation, Software, and Methodology. **Zachary Adam**: Conceptualization, Writing-Review and Editing, Data Curation, **Daniel Cove**: Data Curation, **Betül Kaçar**: Conceptualization, Writing-Review and Editing, Project Administration and Funding acquisition. All authors gave final approval for publication and agreed to be held accountable for the work performed therein.

## Data accessibility

The data is available at the website https://chemorigins.bact.wisc.edu. The code that runs the database is available at the Github repository https://github.com/brunocuevas/chemorigins, and the specific release of this publication at the Zenodo repository https://doi.org/10.5281/zenodo.16792967. The annotations are available at Github repository https://github.com/brunocuevas/chemorigins-data. The specific data release of this publication can be found at https://doi.org/10.5281/zenodo.16792991.

