## Supplemental Information for "ChemOrigins: A community curated database for origins of life chemistry"

### Supplementary figures

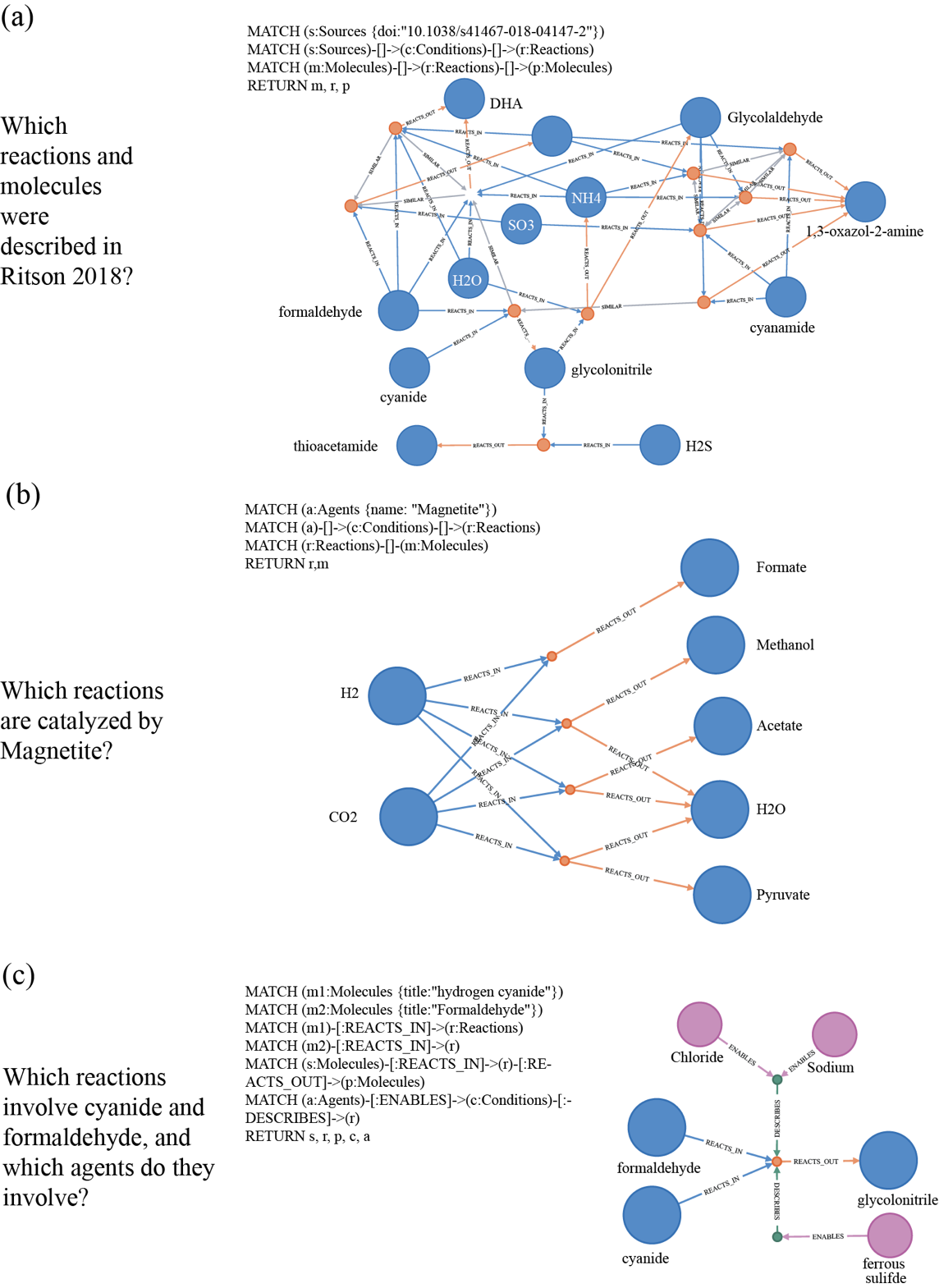

**Figure S1.** Examples of Cypher-based queries using ChemOrigins Neo4J graph database and the results.
